# OICNet: A Neural Network for Online EEG Source Separation using Independent Component Analysis

**DOI:** 10.1101/2023.05.29.542778

**Authors:** Po-Ting Yeh, Arthur C. Tsai, Chia-Ying Hsieh, Chia-Cheng Yang, Chun-Shu Wei

**Affiliations:** Department of Computer Science, National Yang Ming Chiao Tung University (NYCU), Hsinchu, Taiwan; Institute of Education and the Institute of Biomedical Engineering, NYCU; Institute of Statistical Science, Academia Sinica, Taiwan

**Keywords:** Independent Component Analysis, Blind Source Separation, Brain-Computer Interface, Electroencephalography, Deep Neural Networks

## Abstract

Online source separation of EEG signals plays a crucial role in understanding and interpreting brain dynamics in real-time applications such as brain-computer interfaces (BCIs). In this paper, we propose OICNet, a novel neural network designed specifically for online EEG source separation using independent component analysis, aiming to address the challenges of real-time computational efficiency and reliable extraction of independent components from EEG data streams. The OICNet is trained on a loss function that integrates non-Gaussianity measurement and an orthogonality constraint to achieve effective decomposition of multi-channel EEG signals. We conducted comprehensive evaluation of OICNet on both task-related and task-free EEG datasets with comparison against conventional and network-based ICA counterparts. The results demonstrate that OICNet outperforms existing methods in terms of accuracy and computational efficiency. Overall, OICNet provides high-efficiency real-time EEG source separation capabilities and paves the way for advancements in deep-learning EEG processing in real-world BCI applications.

## I. Introduction

**I**ndependent component analysis (ICA) is a signal processing approach that seeks mutually statistically independent sources of an observed multivariate signal [1], [2]. One of the derived applications of ICA that has been widely used is blind source separation (BSS), which in general refers to the task of separating a given mixture of signals into the source signals under the condition of no prior information about the source signals or the mixing process. In biomedical signal processing, ICA has gained great success in the analysis of BSS for biomedical signals such as multi-channel electroencephalography (EEG) recordings [3]–[5]. EEG is one of the most widely used modality for real-time brain computer interface (BCI) because of its non-invasiveness, portability, and high temporal resolution [6]. EEG serves as a powerful tool for tracking changes of brain state such as stress, resting and sleeping based on its high temporal resolution [7], [8]. Nonetheless, EEG recordings contain not only the signal originated from the cortical brain but also artifacts such as eye movements, eye blinking, muscular activity, line noise, etc. The presence of artifacts causes waveform distortion and loss of information in EEG recordings and significantly affects on further analyses and pattern recognition [9].

BSS technique advances the analysis of EEG data by its capability of separating the EEG signal and artifacts, and capturing the nonstatioarity of EEG data [3]. One common use of BSS in EEG analysis is to decompose EEG signals into individual components related to specific patterns such as oscillatory rhythms or event-related potentials (ERPs) [3], [10]. Meanwhile, information of spatial distribution of EEG patterns can be extracted via the processing of BSS. Among the BSS methods for EEG analysis, ICA has been one of the most common tools for isolating various artifacts from contaminated EEG recording as well as extracting activities of brain sources [11].

In order to track the varying brain dynamics, an online ICA algorithm capable for online processing with comparable performance to offline counterparts is highly in demand. Several algorithms have been proposed to perform online ICA in recent decades [12]–[In 2012, Akhtar et al. [15] proposed a recursive ICA algorithm, which was later on applied to online source separation for EEG data and coined as online recursive ICA (ORICA) [17], [18]. However, most ICA algorithms are established on simple linear mixing model which may lead to indistinct source signal extraction due to the nonlinear signal mixture in reality [19], [20]. Meanwhile, the use of closed-form (analytical) solution in the update formula cannot handle the nonlinear source separation because the solution is not unique in nonlinear cases [21].

Recently, novel neural network-based ICA models have been developed to minimize mutual information without the prior knowledge of data such as joint and marginal distribution of data. Furthermore, the model architecture of neural network-based ICA is not constrained by the assumption of linear mixture of source components. Essential traits such as hierarchical layer structure and gradient descent of network-based ICA enable highly flexible design of objective function and training process. Therefore, network-ICA models can support learning of both linear and nonlinear unmixing process. However, existing neural network-based ICA models are often with excessive hyperparameters that pose a challenge in model training and computational efficiency [22]–[24].

Therefore, a novel extensible approach that is capable of tackling the aforementioned issues is needed. We hereby propose an online ICA network (OICNet) dedicated to transforming a set of signal mixture into separated independent components. The proposed OICNet incorporates a composite loss function based on a log-hyperbolic cosine function and a reconstruction term. A task-related and a resting EEG datasets are incorporated to evaluate our method, with comparison against leading conventional and network-based ICA methods. The major contribution of this work is three-fold:

1. We propose a novel neural network-based ICA, OICNet, with a composite loss function for online EEG source separation.
2. We introduce a new metric, Area Under Correlation Evolution (AUCE), designed for assessing the accuracy and efficiency of online BSS algorithms.
3. The proposed OICNet outperforms leading ICA approaches in the efficiency of online EEG source separation and the proximity to the decomposition of the offline ground truth.

## II. Related Work

### A. Conventional Online BSS methods

This section summarizes several representative online BSS methods based on traditional machine learning techniques developed in recent years. Many classic BSS algorithms such as Infomax [25] and FastICA [19] have been widely adopted in EEG source separation. However, their applications are limited to offline analysis since they need the entire dataset or batches of data for computation. In general, the online BSS methods can be divided into two types by their learning algorithms: least-mean-square (LMS) and recursive-least-square (RLS). LMS-type algorithm refers to the adaptation of desired filters with gradient-based approaches and has a low computation cost but a slow convergence speed. EASI [12] and natural-gradient (NG) [13] are two classical methods in this category. In the RLS-type algorithm, the adaptive filter is obtained in a recursive procedure with an exponentially weighted decay based on certain characteristics of the input data. Compared to LMS-type algorithms, RLS-type algorithms mostly feature a faster convergence speed at the expense of computational complexity. A more recent example is ORICA [15], an online ICA algorithm derived from a natural gradient algorithm [13] with a block-update rule. ORICA has been implemented in REST [26], an EEGLAB plugin [27], for online EEG source separation.

### B. Neural Network-based ICA

Recent advances in deep learning techniques have led to the emergence of neural network-based independent component analysis (ICA) approaches. Brakel et al. [22] suggested to minimize JS-divergence, which is an alternative to mutual information, and the reconstruction penalty via adversarial autoencoder. Hlynsson et al. [23] adopted the mutual information neural estimator (MINE) [28], which approximates the lower bound of mutual information as the mutual information estimator. However, these methods require excessive hyperparameter tuning in practice. Recently, Li et al. [24] proposed DDICA that can be trained with non-parametrical mutual information estimator. The mutual information of the independent components (ICs) is computed by the matrix-based Rényi’s *α*-order entropy function. These neural network-based ICA can be considered as an unmixing function and optimized by stochastic gradient descent (SGD) or its variants.

## III. Preliminary

### A. Blind source separation using independent component analysis

In a BSS problem, the measured signal mixtures tend to present Gaussian density distributions, while the source signals tend to retain non-Gaussian density distributions. ICA performs a process of maximizing non-Gaussianity or minimizing mutual information and thus can be applied to unmixing signal mixtures. The category of ICA algorithms that aim to minimize mutual information measures the dependence between estimated sources to extract the sources from a given mixture. However, the density of observed signal is often difficult to estimate with limited data in practice, which poses a challenge to compute mutual information efficiently. The computation of mutual information needs the distribution assumption or complex approximation and thus is costly to estimate practically. For the non-Gaussianity maximization counterpart, Comon suggests using negentropy as a measurement of non-Gaussianity of each estimated sources [2]. With differential entropy *H* in information theory, the randomness of a given variable can be assessed. One important property of entropy is that it is the largest when the given variable follows Gaussian distribution. The differential entropy *H* for a random vector **y** with density *p*(*·*) is defined as:

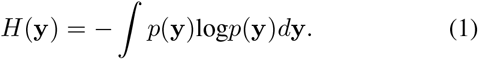

Unfortunately, differential entropy is not invariant for any invertible transformation. Comon have shown that negentropy is always positive and invariant for linear transformation [2], and negentropy *J* is defined as follows:

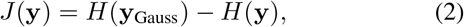

where **y**_Gauss_ denotes a Gaussian variable with the same covariance matrix with **y**. After that, Hyvarinen further proposes a new approximation of negentropy with some smooth contrast functions *G* to simplify the computation [29], [30]:

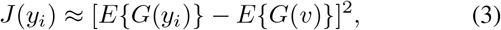

where *v* is a normalized Gaussian variable.

A common assumption in the ICA problem is that the observed mixture is a linear combination of underlying sources:

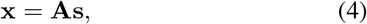

where the vector **x** = [*x*_1_, *x*_2_, …, *x*_*n*_] ∈ ℝ^*n*^ is the observed *n*-dimensional signal, the vector **s** = [*s*_1_, *s*_2_, …, *s*_*m*_] ∈ℝ^*m*^ is the *m*-dimensional statistically independent components and **A** ∈ℝ^*n×m*^ denotes the mixing matrix. The objective of the ICA algorithm is to determine the inverse of mixing matrix **W** = **A**^*−*1^, which decomposes the observed data:

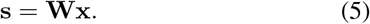

Furthermore, most studies assume that the number of observed variables is equal to the number of ICs, i.e. *n* = *m*.

### B. Prewhitening

The problem of ICA can be simplified by whitening the observed data **x** = [*x*_1_, *x*_2_, …, *x*_*n*_] ∈ℝ^*n*^ before performing ICA, as this reduces the number of parameters [31]. The idea behind whitening is to transform the data **x** through a linear transformation **U** *∈* ℝ^*n×n*^ into a new representation:

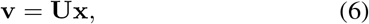

where the components of transformed data **v** = [*v*_1_, *v*_2_, …, *v*_*n*_] ∈ℝ^*n*^ are uncorrelated and have unit-variance, i.e., the covariance matrix of the **v** is equivalent to the identity matrix *E* **vv**^T^ = **I**. For example, a widely used method is based on the eigenvalue decomposition (EVD) of the covariance matrix **C** = *E* **xx**^T^ = **EDE**^T^, where **E** is an orthogonal matrix of eigenvectors of the covariance matrix **C** and **D** is a diagonal matrix that contains the eigenvalues corresponding to **E**. A whitening matrix **U** is therefore built by **U** = **ED**^*−*1*/*2^**E**^T^ where **D**^*−*1*/*2^ is element-wise inverse square root of **D**. With the whitening matrix **U**, the ICA model is expressed as:

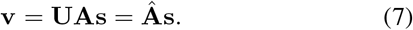

This whitening transformation imposes an orthogonal constraint on the mixing matrix **Â**:

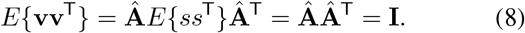

According to Eq. 8, the number of parameters required for estimation the mixing matrix is reduced since the degree of freedom is reduced from its original value *n*^2^ to *n*(*n −*1)*/*2 due to the orthogonal constraint [31].

## IV. Online Independent Component Analysis Network

### A. Model Architecture

The architecture of the proposed OICNet is illustrated as in Fig. 1. In a nutshell, the autoencoder-like architecture comprises a single-layer encoder and a single layer decoder, where the encoder (former layer), parameterized by **W**, transforms a segment of multi-variate time series data into corresponding ICs, and a decoder (latter layer), parameterized by **W**^T^, reconstructs the data via back projection of the ICs onto the original domain. Note that the weights of decoder are tied with encoder to reduce the number of parameter of the model. In the case of EEG decomposition, a segment of multichannel EEG time series **X** ∈ℝ^*C×T*^ with *C* electrodes and *T* time points, the ICs are extracted by *C* spatial kernels with size (*C*, 1) in this work. Although the number of ICs in this model can be any positive integers, it was set to be the same as the number of channels *C* by default. This was done to enable comparative validation against other baseline methods that assume the number of sources is equal to the number of channels.

**Figure 1.**
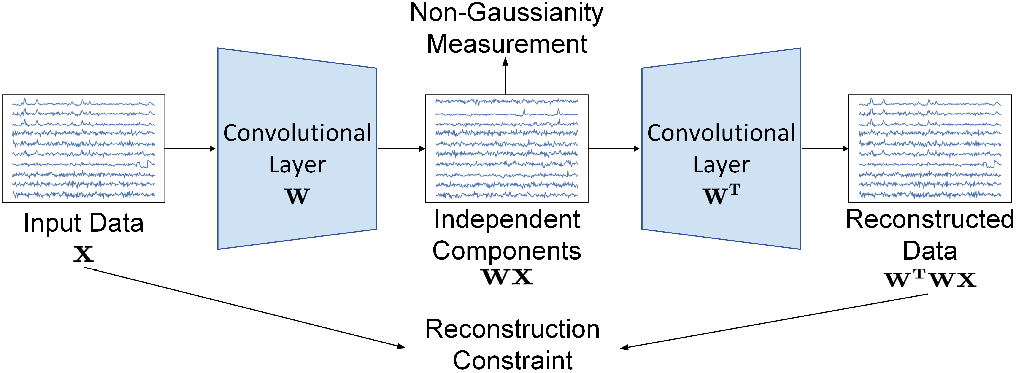
An overview of OICNet’s architecture. The OICNet consists of a single convolutional layer that performs unmixing projection for the input channel-domain data **X**. The loss function comprises 1) a non-Gaussianity measurement that is qualified by a contrast function using the unmixed independent components **WX** and 2) a reconstruction constraint that assures the orthogonality of the unmixing matrix **W** using the back projected reconstructed data **W**^T^**WX**.

### B. Loss Function

This section presents the design of the loss function used to fine-tune OICNet. The loss function incorporates two key components, a measurement of the non-Gaussianity of independent components and a reconstruction constraint applied to kernels, as expressed in the equation:

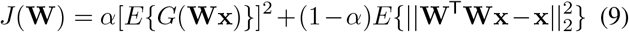

where *G* is practically any nonquadratic function or contrast function [29], || *·* ||_2_ denotes *L*_2_ norm. As discussed in the previous section, the contrast function is used to measure the non-Gaussianity of a random variable. We exploit the hyperbolic cosine function and pass the estimated sources through a tanh(*·*) activation function before computing non-Gaussianity [30]:

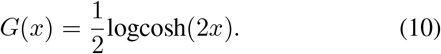

The reconstruction constraint is another crucial component of the loss function. It is designed to prevent the replication of features and enhance the robustness of OICNet to incompletely whitened data. The reconstruction constraint improves the performance compared to the orthogonality constraint in the standard ICA problem [32]. Le et al. [32] have shown that if the data is whitened, the reconstruction constraint is equivalent to the orthogonality constraint. Orthogonality prohibits kernels from producing similar features and preserves the whiteness of independent components, and experimental results suggest that the reconstruction constraint is less sensitive to whitening, indicating that the ICA model can still perform well even when the input data is insufficient [33].

### C. Prewhitening

As previously mentioned, whitening the data is a crucial step that facilitates the estimation of the unmixing matrix **W** [31]. To this end, we implemented an online RLS-type prewhitening algorithm in our framework [34]:

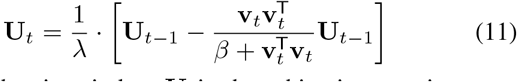

where *t* is the time index, **U** is the whitening matrix, **v**_*t*_ = **U**_*t−*1_**x**_*t*_ is the whitened data at time *t, λ*(0 *< λ ≤*1) is a forgetting factor to determine how fast the algorithm neglects past data, and its associated coefficient *β* is defined as *λ/*(1*− λ*). Once a certain amount of data **X** = [**x**_1_, **x**_2_, …, **x**_*n*_] *∈* ℝ^*m×n*^ was collected, we updated the whitening matrix **U** using a nonoverlapping block of data **x**_*t*_ with a block size *K*_*b*_. This approach minimizes computational overhead while ensuring that the data is sufficiently prewhitened for the subsequent unmixing process [18].

### D. Online Model Tuning

The proposed online ICA approach includes an online whitening method and OICNet, the framework is depicted in Fig. 2 with the pseudo codes are present in Algorithm 1. To begin the process, a batch of data was given, and both the whitening matrix **U** and the unmixing matrix **W** were initialized as identity matrices. The whitening matrix **U** was updated using the Eq. 11. Next, the batch of data was processed by multiplying the whitening matrix **U** to acquire the prewhitened data. This procedure ensured that the data was prewhitened before being used to fine-tune the model, thereby enhancing the stability of the OICNet. Next, the prewhitened data were used to update the unmixing matrix **W** using Eq. 9 with the Adam optimizer [35]. It is important to emphasize that the entire model tuning process for one batch of data is expected to be completed prior to receiving the next batch of data in the online pipeline. We conducted a grid search to determine the optimal parameter configurations of *η, α, K*_*b*_, and the number of epochs *E* for each fine-tuning. After experimentation, we found that a block size of *K*_*b*_ = 16 was optimal for the RLS whitening procedure, consistent with previous findings in [18], and was used for the OICNet.

**Figure 2.**
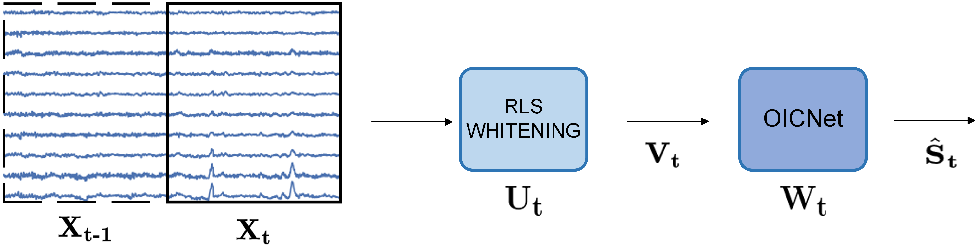
The online processing flow of the OICNet with RLS prewhitening. The EEG data **X** is sliced by a moving time window. At the *t*-th update, the EEG epoch is prewhitened to obtain **V**_*t*_, and then is unmixed by the unmixing matrix of OICNet **W**_*t*_to output the estimated source activity **Ŝ**_*t*_.

## V. Experiments

### A. Data and Preprocessing

This study aimed to evaluate the performance of an online BSS method using different types of data. We employed a set of synthetic data, generated by simulating a mixture of synthetic ground truth signals, to validate the ability of the proposed OICNet to reconstruct sources accurately. Furthermore, we incorporated two real EEG datasets to quantify the performance of online BSS methods.

#### Synthetic Data - Multivariate Time Series

To validate the proposed OICNet’s performance in blind source separation, we artificially synthesized a random mixture of a set of eight source time series. The eight source signals were generated based on a modified synthetic source signals used in the previous study [15]. Specifically, *s*_1_ consists of four peaks to simulate eye blinks, *s*_2_ and *s*_3_ are sinusoidal waves of 10Hz and 25Hz to simulate alpha and beta rhythm, respectively. *s*_4_ is the random signal generated by a uniform distribution on the interval [–1, 1) using the *Numpy* function *random*.*uniform*. The remaining signals are generated by following equations: *s*_5_ = sin(400*t*)cos(30*t*), *s*_6_ = sign(sin(550*t*) + 9cos(40*t*)), *s*_7_ = cos(400*t* + 10sin(90*t*)) and *s*_8_ = sign(cos(550*t*) 5sin(99*t*)). Furthermore, a random noise sampled from the interval [0, 0.05) uniformly was applied to each of the source signals. Finally, an 8 × 8 mixing matrix **A** was generated randomly by uniform distribution on the interval [–1, 1) to project the source signals into a set of multivariate time series signals that simulates the mixture of sources.

##### Algorithm 1

Training algorithms of OICNet on the prewhitening matrix **U** and the unmixing matrix **W**

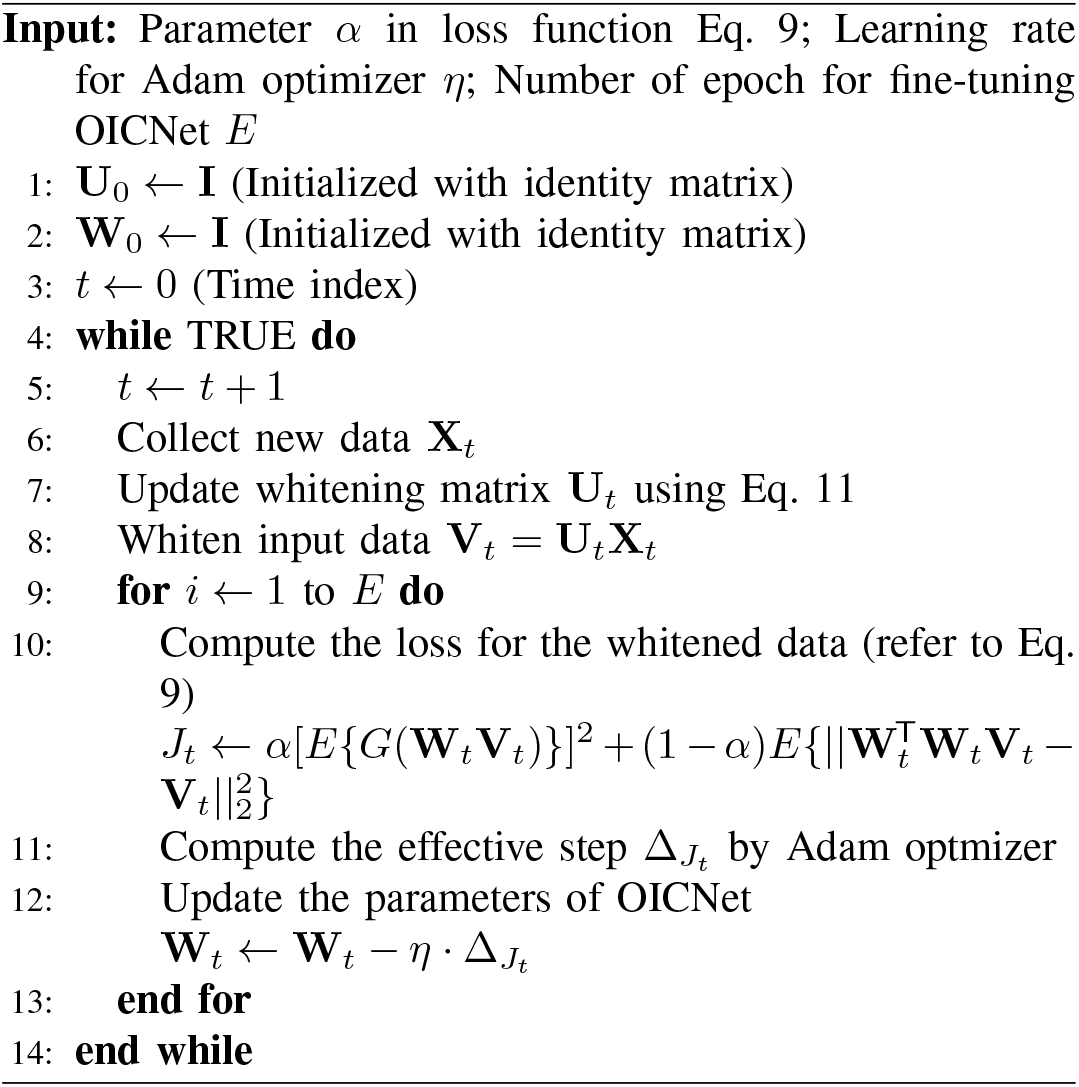

#### EEG Dataset I - Event-related potential (ERP) EEG

This dataset is known as the Kaggle BCI Challenge dataset and includes EEG recordings from 26 subjects performing a P300-based BCI spelling task [36]. The objective of the experiment was to identify and classify the signal perturbations that arose from erroneous feedback in order to improve the overall robustness of the BCI speller. EEG recordings were made using a 56 Ag/AgCl electrode arranged in accordance with the extended 10-20 system and were sampled at a rate of 600 Hz. Each subject participated in five sessions, with the first four sessions comprising 60 trials and the last session comprising 100 trials. There were three types of ERPs waveforms to be extracted from the EEG recordings. The first type of ERP was measured in response to the flashing of visual stimuli as a visual evoked potential (VEP). The other two types of ERP were measured in response to the correct/erroneous feedback from the recognition results of the BCI speller as distinguishable ERP waveforms. Specifically, the ERP waveforms corresponding to the erroneous feedbacks were termed error-related potential (ErrP) or error-related negativity (ERN) in literature [37]. For the purposes of the study, the 16 subjects who were released early in the Kaggle competition were used [38]. Preprocessing steps included downsampling from 600 Hz to 128 Hz and band-pass filtering at 1-40 Hz.

#### EEG Dataset II - Resting eye-open/closed EEG

We recruited ten healthy participants and recorded their restingstate EEG using a 30 Ag/AgCl electrodes EEG cap at a 1K sampling rate. This experimental protocol was approved by the Institutional Review Board of National Yang Ming Chiao Tung University, Hsinchu, Taiwan. All the subjects were asked to read and sign an informed consent form prior to the experiments. Each recording lasted for one minute and comprised a session of 30-second eye-closed resting state followed by a session of 30-second eye-open resting state. The EEG signal was downsampled to 250 Hz, applying common average re-reference and high-passing filtering with a cutoff of 1 Hz.

### B. Performance Evaluation

In this work, we determine the performance of an online ICA method by evaluating how similar the resulting source separation is to the ground truth offered by a classic offline method on real EEG data. Since the true unmixing matrix is not available for real EEG data, one approach to quantitative evaluation for online ICA is to compare with an reliable ICA method performed offline [17]. To achieve this, we employed the widely used Infomax ICA to generate the ground truth of unmixing matrices. Additionally, we proposed a specific metric for assessing the similarity between an evolving component given by an online method and its corresponding static component given by the Infomax ICA.

#### 1) Ground Truth of Source Separation

Infomax ICA has been considered as one of the most widely used and effective offline ICA methods according to [39]. In order to compare the performance of different ICA methods, we need a reliable ground truth for evaluation. Therefore, in this study, we used the results obtained from the Infomax function offered by MNE-python [40] as the “gold standard” ground truth for comparison. We first identified a list of best-matching components between the ground truth Infomax ICA and the decomposition results obtained from other ICA methods. This was done by a component matching process as described in the following paragraph. After the best-matching components were identified, we assessed the similarity between the two sets of components and compare the performance of different ICA methods. However, it should be noted that the results of EEG source separation can vary across ICA methods that are evaluated.

#### 2) Component Matching

To monitor the converging evolution of an IC during online tuning, a method for matching ICs is required. We match the spatial pattern of one online algorithm and the offline ground truth offered by the Infomax ICA, which is a column in the mixing matrix, and estimate their similarity based on the approach proposed in [41]:

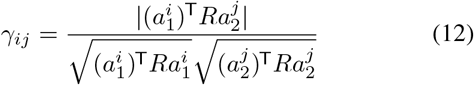

##### Algorithm 2

Matching algorithm for components from two ICA methods

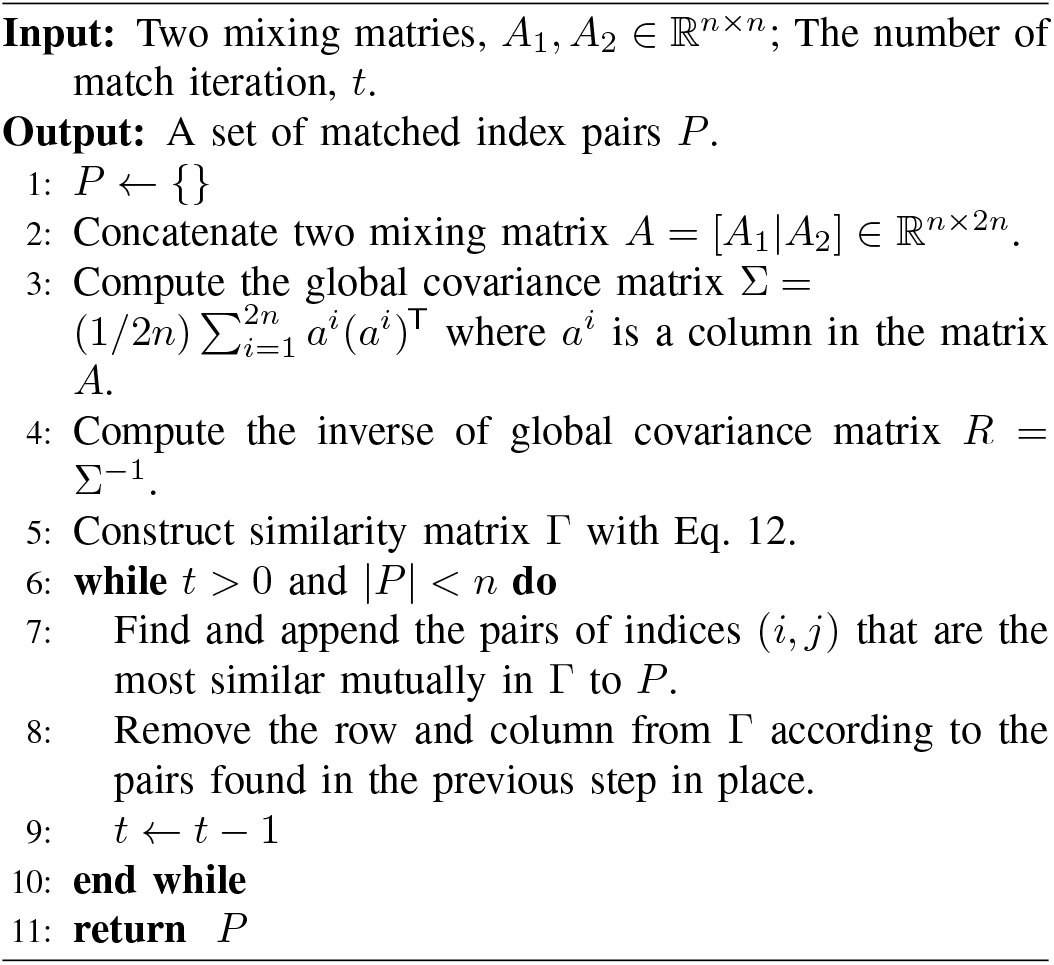

where 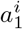 and 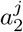 are columns from mixing matrice *A*_1_ and *A*_2_, *R* is the inverse of the “global” covariance 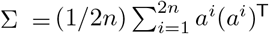 where *a*^*i*^ is a column in the matrix *A* = [*A*_1_|*A*_2_] ∈ℝ^*n×*2*n*^. The matching process involves identifying pairs of components from different methods that are most similar to each other. To facilitate this process, we developed a matching procedure that sequentially matches the unmatched components until a desired criterion is met. In this study, we defined a common component as a ground truth independent component that can be matched by all methods. We continue the matching process until a certain criterion, such as the percentage of variance explained by the common components being at least 80%, is satisfied. This ensures a sufficient variance explained by the common components matched with the ground truth. The matching procedure is further detailed in Algorithm 2.

#### 3) Component Similarity

With the ground truth Infomax ICA method and an online ICA method to be evaluated, we can trace the evolving process of an IC in an online model tuning process. To quantify the convergence speed and stability of an online ICA algorithm, we define a metric called Area Under Correlation Evolution curve (AUCE). AUCE is defined as the average correlation between matched components throughout the duration of online tuning. Given a component **Ŝ** obtained from an online method and **S** is its corresponding ground truth, the AUCE of this pair is calculated as:

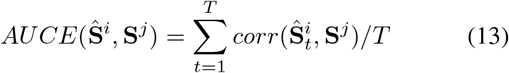

where *corr*(*·*) denotes the magnitude of Pearson’s correlation, *i* and *j* are row indices, *t* represents the index of update, and *T* is the total number of time indices. The AUCE ranges from (12) 0 to 1 and is affected by the speed of convergence and the similarity between matched components.

## VI. Results

This section describes the results obtained from the experiments using the synthetic data and two datasets of real EEG. The performance of the proposed OICNet was validated and compared against other ICA methods in a simulated online BSS scenario.

### A. Validation on Synthetic Data

To demonstrate the capability of OICNet for online BSS with corrective updates, we compared the source signals, mixed signals, and reconstructed source signals using Fig. 3. Fig. 3(a) displays the ground truth synthetic source signals, while Fig. 3(b) shows the mixed signals. Fig. 3(c), on the other hand, illustrates the reconstructed source signals. The reconstructed source signals are visually indistinguishable from the ground truth signals, indicating OICNet’s ability to perform accurate blind source separation.

**Figure 3.**
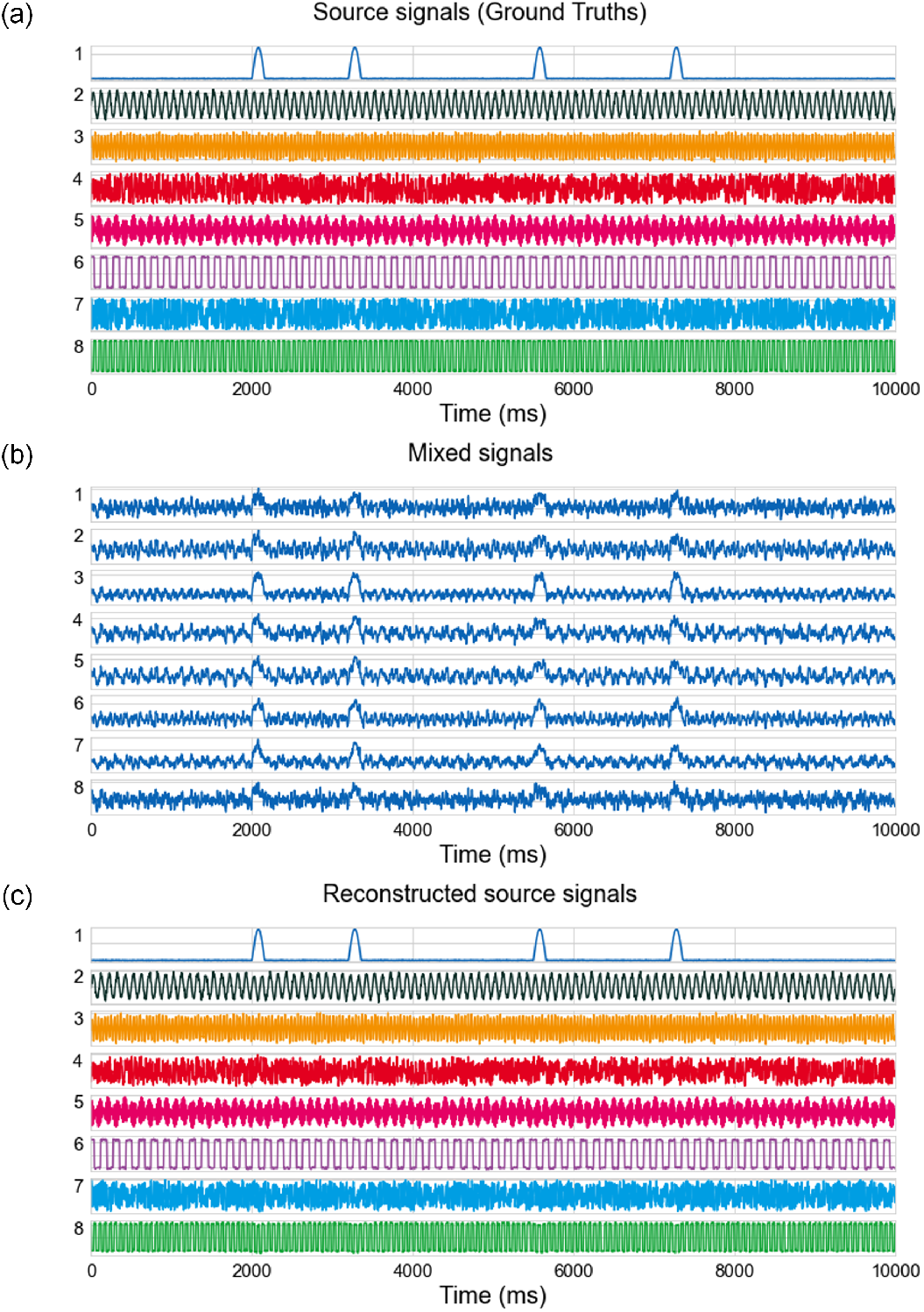
The waveforms of simulated data. (a) The sources, (b) the mixed signals and (c) the sources reconstructed by OICNet. The reconstructed signals by the OICNet exhibited a high degree of similarity to the sources signal. This primary experiment provided the compelling validation of the OICNet’s ability for online source separation.

### B. Validation on Real EEG Data

We applied the proposed OICNet to online source separation on real EEG data accessed from the ERP and the resting EEG datasets. The Infomax ICA was employed as the ground truth for comparing the performance of OICNet in extracting ERP and artifact waveforms from the Kaggle BCI Challenge dataset, as well as capturing the dynamic of EEG components in the transitioning between eye-closed and eye-open state from the resting EEG dataset.

### 1) Source Separation on ERP Data

To validate the capability of OICNet for separating the sources of ERP responses extracted from the Kaggle BCI Challenge dataset, Fig. 4 compares the EEG patterns. The top row shows the ERN, VEP, and eye-blink waveforms extracted by Infomax ICA, while the bottom row shows those extracted by OICNet. Fig. 4(a) depicts the ERN waveforms elicited by incorrect and correct feedback of the BCI speller [37]. Both Infomax ICA and OICNet were able to separate the ERN component at the frontocentral area. Fig. 4(b) shows the VEP waveforms consisting of N70, P1, and N140 ERP components originating from the parietal and occipital lobes [42]. Fig. 4(c) illustrates a single eye-blink artifact that severely contaminates frontal channels with a strong burst in raw EEG data. We observed highly analogous spatial distributions and temporal patterns of the selected features obtained from both methods. These results demonstrate that OICNet is capable of isolating informative brain activities and artifacts.

**Figure 4.**
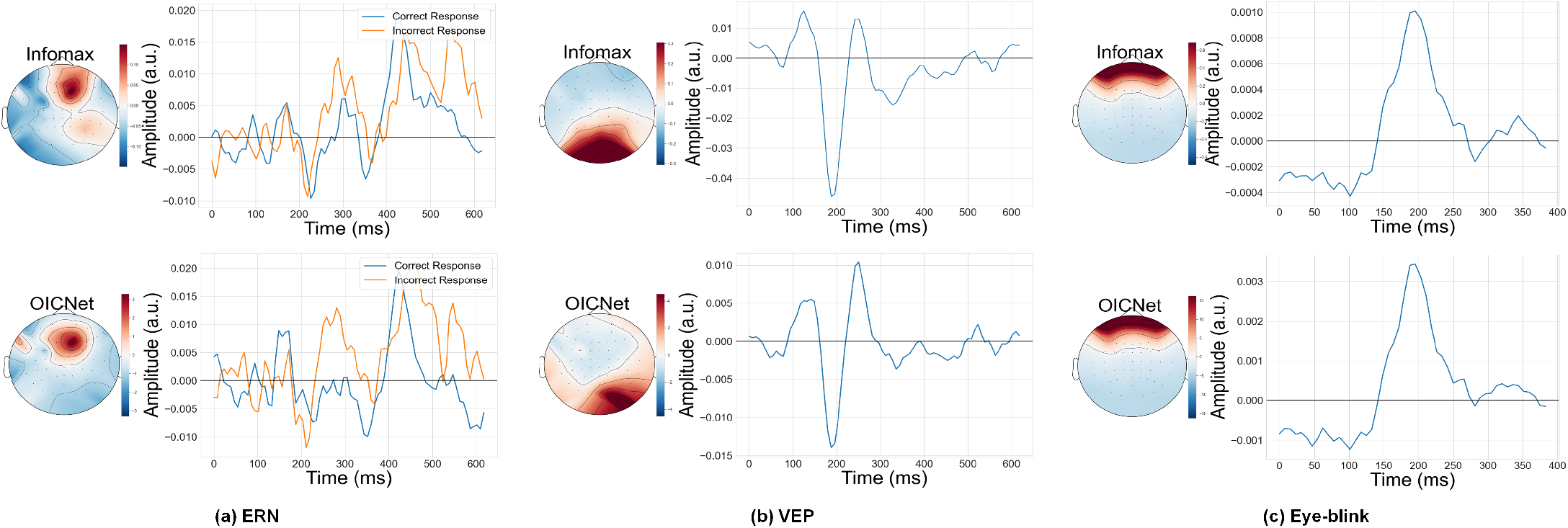
Waveforms of (a) ERN, (b) VEP, and (c) eye-blink artifact extracted by Infomax ICA (top) and the proposed OICNet (bottom) from S17 of the ERP dataset. (a) illustrates the responses to incorrect/correct feedback of a BCI spller marked in red/blue. (b) illustrates VEP waveforms. (c) illustrates a single eye-blink artifact.

### 2) Source Separation on Resting EEG Data

Fig. 5 displays the correlation between the components isolated by OICNet and their corresponding Infomax ICA ground truths. We selected two OICNet components that match the eye-blink and occipital alpha components, respectively, extracted by Infomax ICA. During the entire 30-second eye-closed session, the OICNet component #30 shows a high correlation with the Infomax ICA occipital alpha component. Immediately after the onset of the 30-second eye-open session, the correlation drops, and the OICNet component #1 displays increasing correlation with the Infomax ICA eye-blink component. These results demonstrate OICNet’s high sensitivity to non-stationary EEG data and its ability to promptly adapt to different brain states.

**Figure 5.**
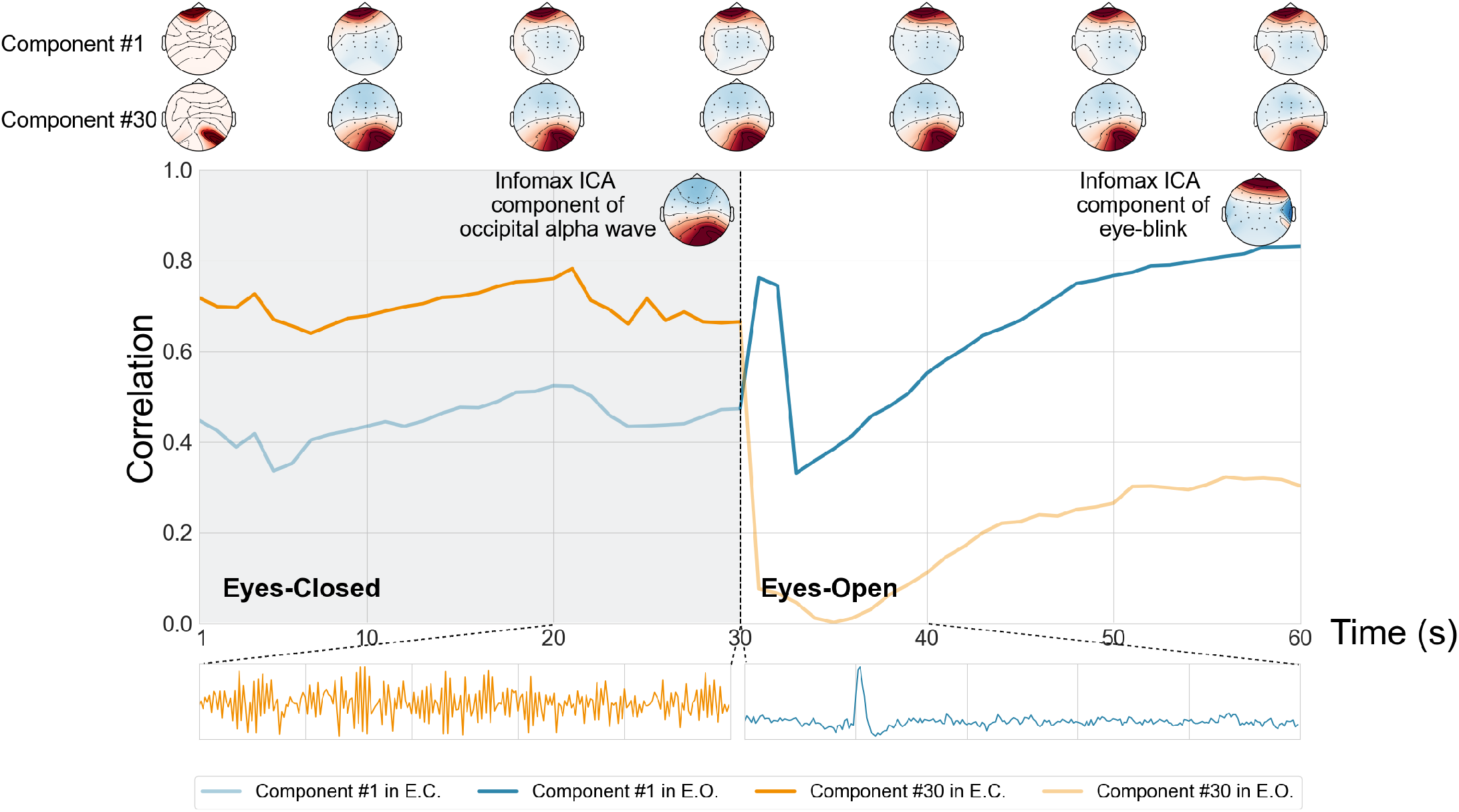
The changes of selected components extracted by OICNet in terms of correlation and topoplot on the private EEG dataset. The entire session lasts sixty seconds and consists of two resting states: eyes-closed state (E.C.) for the first thirty seconds and eyes-open state (E.O.) for the remaining time. The transformations of eye-blink artifact (component #1) and alpha wave (component #30) are illustrated at the top of the figure and their waveforms are depicted below. The vertical dotted line presents the time of state change and the ground truths offered by Infomax ICA are shown in the upper right of each session.

### C. Performance Comparison

We compare its performance with several other online ICA methods, including ORICA [18], online implementations of ANICA [22], MINE [23], and DDICA [24]. We present quantitative results to further validate the efficiency of OICNet in real-time source separation tasks. This comprehensive evaluation provides a solid basis for assessing the performance of OICNet and its superiority over other online ICA methods. As shown in Fig. 6, the mean correlation curve of OICNet exhibits a rapid increase at the beginning of the session, followed by a high plateau that is maintained until the end. Table I summarizes the results, showing that OICNet provides the highest AUCE across all subjects as well as within individual subjects.

**Figure 6.**
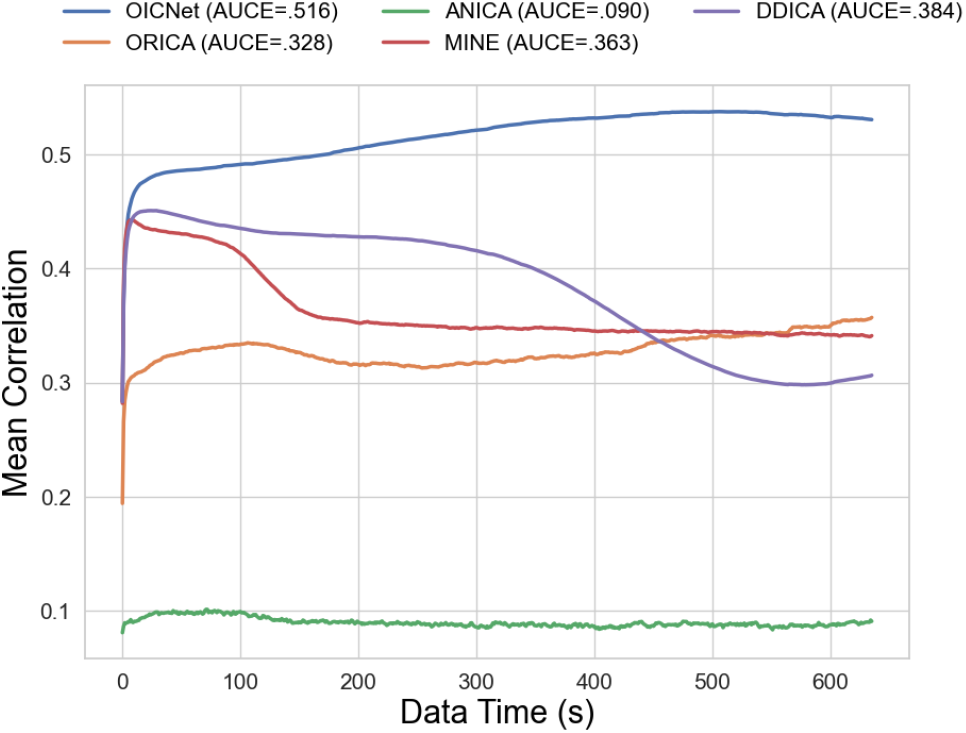
The comparison of mean correlation evolution of common components over all sessions from the training set in the ERP dataset. The timeline was truncated to fit the shortest session for the evaluation of average performance. Among the methods evaluated, the OICNet exhibited the best tuning stability and the resulting independent components were the most similar to the ground truth. The metric AUCE is computed as the area between the correlation curve and the x-axis as in Eq. 13.

**Table I.**
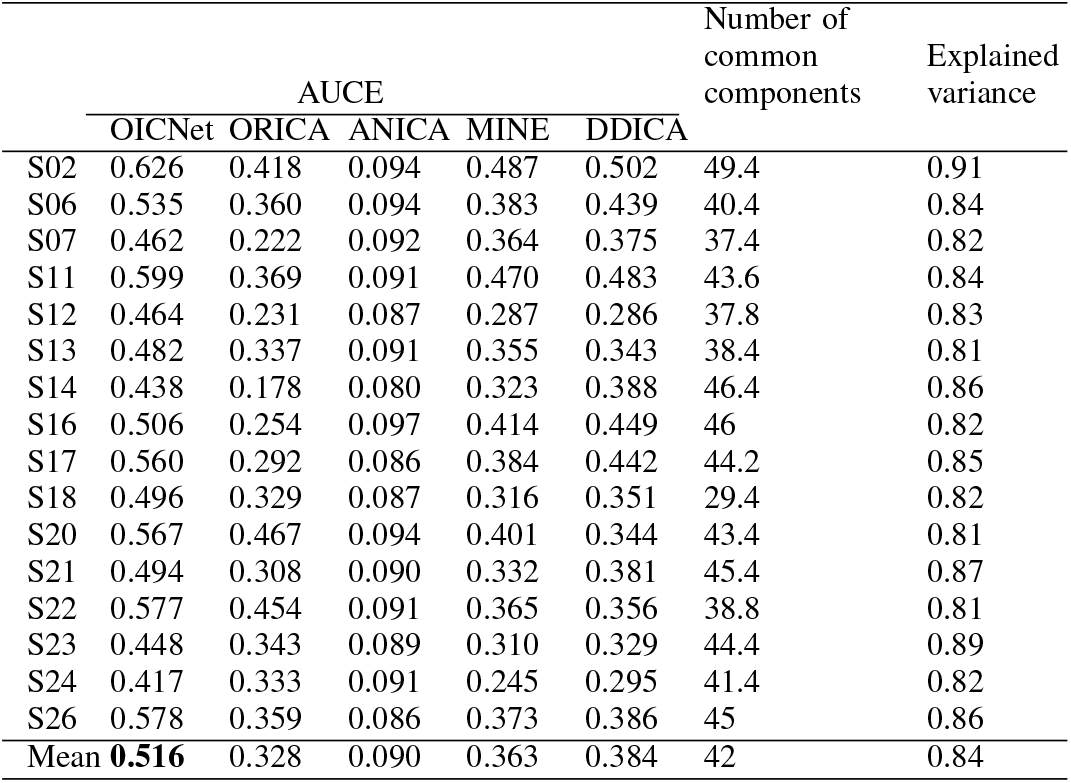
Performance comparison on the ERP EEG Data.

|  | AUCE |  |  |  |  | Number of common components | Explained variance |
| --- | --- | --- | --- | --- | --- | --- | --- |
|  | OICNet | ORICA | ANICA | MINE | DDICA |  |  |
| S02 | 0.626 | 0.418 | 0.094 | 0.487 | 0.502 | 49.4 | 0.91 |
| S06 | 0.535 | 0.360 | 0.094 | 0.383 | 0.439 | 40.4 | 0.84 |
| S07 | 0.462 | 0.222 | 0.092 | 0.364 | 0.375 | 37.4 | 0.82 |
| S11 | 0.599 | 0.369 | 0.091 | 0.470 | 0.483 | 43.6 | 0.84 |
| S12 | 0.464 | 0.231 | 0.087 | 0.287 | 0.286 | 37.8 | 0.83 |
| S13 | 0.482 | 0.337 | 0.091 | 0.355 | 0.343 | 38.4 | 0.81 |
| S14 | 0.438 | 0.178 | 0.080 | 0.323 | 0.388 | 46.4 | 0.86 |
| S16 | 0.506 | 0.254 | 0.097 | 0.414 | 0.449 | 46 | 0.82 |
| S17 | 0.560 | 0.292 | 0.086 | 0.384 | 0.442 | 44.2 | 0.85 |
| S18 | 0.496 | 0.329 | 0.087 | 0.316 | 0.351 | 29.4 | 0.82 |
| S20 | 0.567 | 0.467 | 0.094 | 0.401 | 0.344 | 43.4 | 0.81 |
| S21 | 0.494 | 0.308 | 0.090 | 0.332 | 0.381 | 45.4 | 0.87 |
| S22 | 0.577 | 0.454 | 0.091 | 0.365 | 0.356 | 38.8 | 0.81 |
| S23 | 0.448 | 0.343 | 0.089 | 0.310 | 0.329 | 44.4 | 0.89 |
| S24 | 0.417 | 0.333 | 0.091 | 0.245 | 0.295 | 41.4 | 0.82 |
| S26 | 0.578 | 0.359 | 0.086 | 0.373 | 0.386 | 45 | 0.86 |
| Mean | **0.516** | 0.328 | 0.090 | 0.363 | 0.384 | 42 | 0.84 |

For the resting EEG dataset, we present the mean correlation evolution curves in Fig. 7. Most online ICA methods exhibit rapid convergence of the mean correlation from the beginning of the session until a jump at 30 seconds in response to the transition between eye-closed and eye-open sessions. OICNet again offers the highest correlation evolution curve above other methods, although the difference is not as apparent as in the case of the ERP EEG dataset. Table II shows that OICNet

**Figure 7.**
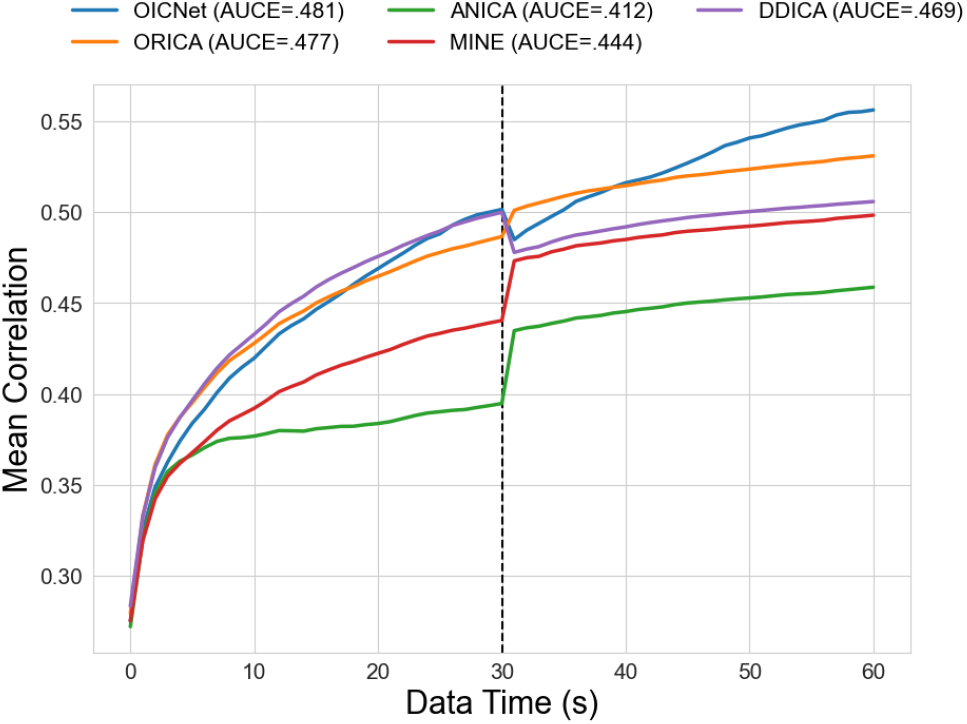
The comparison of mean correlation evolution of common components for the resting EEG dataset. The vertical dotted line indicated that the state switch from eye-closed to eye-open. Remarkably, the OICNet demonstrated the highest correlation evolution curve after the state changed, suggesting its superior adaptivity to different state compared to other methods.

**Table II.** Performance comparison on the resting EEG Data.

|  | Average AUCE |  |  |  |  | Average number of common components | Average explained variance |
| --- | --- | --- | --- | --- | --- | --- | --- |
|  | OICNet | ORICA | ANICA | MINE | DDICA |  |  |
| S01 | 0.498 | 0.480 | 0.411 | 0.443 | 0.491 | 24 | 0.83 |
| S02 | 0.510 | 0.519 | 0.464 | 0.488 | 0.515 | 24.5 | 0.82 |
| S03 | 0.449 | 0.414 | 0.342 | 0.359 | 0.318 | 24 | 0.84 |
| S04 | 0.509 | 0.502 | 0.425 | 0.465 | 0.500 | 24.5 | 0.86 |
| S05 | 0.421 | 0.427 | 0.379 | 0.383 | 0.404 | 24.5 | 0.83 |
| S06 | 0.518 | 0.532 | 0.472 | 0.486 | 0.520 | 23 | 0.82 |
| S07 | 0.416 | 0.446 | 0.384 | 0.436 | 0.454 | 22 | 0.81 |
| S08 | 0.527 | 0.538 | 0.468 | 0.505 | 0.534 | 22.5 | 0.82 |
| S09 | 0.445 | 0.407 | 0.334 | 0.385 | 0.425 | 25 | 0.86 |
| S10 | 0.519 | 0.507 | 0.438 | 0.487 | 0.533 | 22.5 | 0.81 |
| Mean | **0.481** | 0.477 | 0.412 | 0.444 | 0.469 | 23.65 | 0.83 |

provides the highest overall AUCE. It is important to note that the online implementation of ANICA underwent additional parameter tuning (number of iterations reduced to 200) due to limited computational power on a local machine. Therefore, the comparison takes this limitation into account.

Moreover, we evaluate the efficiency of OICNet in comparison with other methods by estimating the average time elapsed for fine-tuning with 250 time points for each method. The results, presented in Table III, revealed that OICNet emerged as the fastest algorithm on both datasets. It demonstrated remarkable computational efficiency, characterized by low processing time and the ability to effectively handle multi-channel EEG data streams in real-time scenarios. This efficiency aligns closely with the ground truth of Infomax ICA, emphasizing the practicality and suitability of OICNet for online EEG source separation tasks.

**Table III.** Mean elapsed time (ms) for processing a single batch of EEG Data (250 Time points) In a fine-tuning epoch.

| Method | OICNet | ORICA | ANICA | MINE | DDICA |
| --- | --- | --- | --- | --- | --- |
| Dataset I<br>(56-ch,<br>1953 ms/batch) | 9.127 | 14.647 | 382.673 | 162.927 | 264.151 |
| Dataset II<br>(30-ch,<br>1000 ms/batch) | 5.812 | 11.543 | 367.736 | 80.054 | 150.528 |

Note: All simulations were conducted on an Intel® Core™ i7-10700F CPU operating at 2.90GHz.

### D. Ablation Studies

In the ablation studies, we varied the weight coefficient *α* from 0 to 1 to determine the optimal combination of the non-Gaussianity measurement and the orthogonality constraint. We evaluated the overall AUCE performance on both dataset. The results, presented in Fig. 8, indicate that the best performance is achieved when *α* = 0.994 for both the ERP and resting EEG datasets. Additionally, comparable performances are observed when *α* is in the range of [0.99, 0.994]. However, when *α* approaches 1, the AUCE performance decreases significantly as the loss function becomes solely dependent on non-Gaussianity measurement.

**Figure 8.**
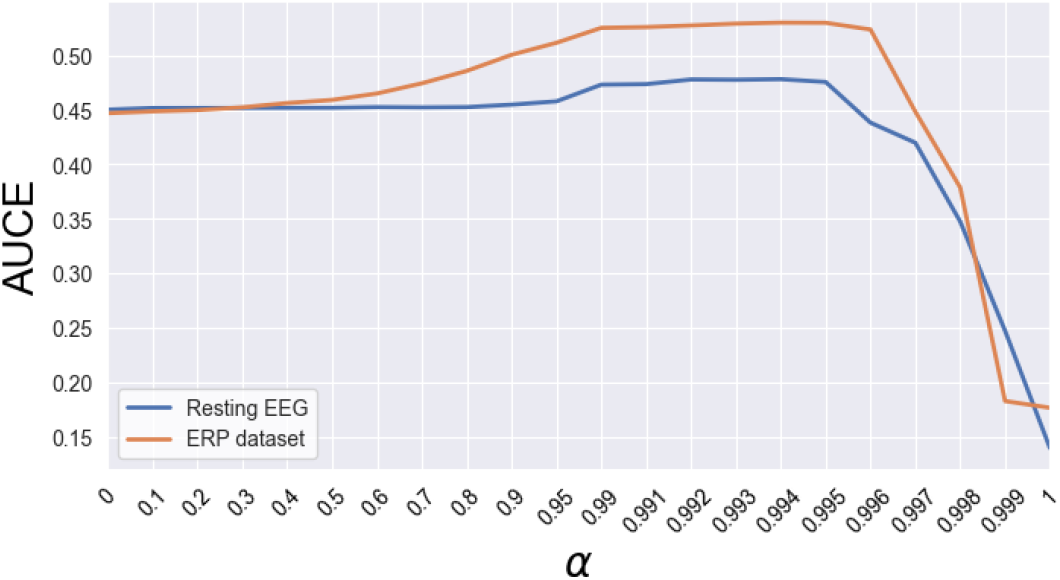
The investigation of different *α* values in the loss function on both datasets revealed that the choice of *α* ranging from 0.99 to 0.994 yielded comparable results, indicating that the weight selection may not be a critical issue for user.

We also conducted experiments to evaluate the performance of different types of non-Gaussianity measurement and orthogonality constraint using the resting EEG data. The results in Table IV demonstrate that the reconstruction constraint in OICNet achieves the highest overall performance compared to other methods such as Bansal et al. [43], mutual coherence (MC), spectral restricted isometry property (SRIP), or no orthogonal regularization constraint (None). Interestingly, the choice of non-Gaussianity measurement appears to be less critical when the reconstruction constraint is employed, as all three contrast functions exhibit comparable performance.

**Table IV.** Ablation analysis on the use of non-Gaussianity measurements and orthogonal regularization methods using the resting EEG data.

| | $\text{logcosh}(x)$ | $-\exp(-x^2/2)$ | $x^4/4$ |
| --- | --- | --- | --- |
| REC | **0.470** | 0.420 | 0.453 |
| MC | 0.140 | 0.132 | 0.143 |
| SRIP | 0.140 | 0.203 | 0.141 |
| None | 0.142 | 0.169 | 0.148 |

## VII. Discussion

### A. Fast and Stable Training Process

Firstly, OICNet is robust to unwhitened data, which is beneficial for rapid source separation in the early stages when the whitening matrix has not yet converged. This reduces the dependence of ICA on whitened data and the system overhead induced by whitening algorithms. Secondly, OICNet exhibits stable learning of independent components, as demonstrated by the steady correlation curve. This indicates that the permutation of ICs does not change arbitrarily and that the components appearing later are based on similar ICs from previous sessions. Additionally, OICNet is computationally efficient, as evidenced by its superior performance in terms of computational cost compared to other methods, as shown in Table III. Notably, ORICA was tested with block sizes of 16 and 8 for RLS whitening and ICA stages, respectively, as recommended in [18]. Compared to DDICA, which also optimizes the unmixing matrix using a loss function, OICNet’s loss function is more computationally efficient for online EEG source separation. It is worth noting that ANICA and MINE involve other modules that can be customized by users, but we used the default settings in their open-source code. Furthermore, ANICA requires a large number of iterations to achieve optimal results [22], making it less suitable for online applications.

### B. Whitening Method and Loss Function

Although RLS whitening has been shown to work well with OICNet, exploring alternative whitening methods may be beneficial. However, the suitability of these methods needs to be validated, and parameter selection must be carefully coordinated to ensure successful integration with OICNet’s learning rule. The unconstrained nature of OICNet’s loss function Eq. 9 allows for easy attachment or combination with other loss functions or regularization techniques to facilitate the training process. For example, the contrast functions can be combined with careful weight assignments for each other to capture more diverse EEG features [30]. The results of our ablation study reveal that the orthogonal constraints, apart from the reconstruction constraint (REC), are not applicable for all contrast functions. This observation can be attributed to the fact that the calculation of REC involves the current data. Surprisingly, when using the pure reconstruction constraint as the loss function for OICNet, where *α* is set to 0, the performance is comparable to when the contrast function is adopted. This result indicates that the reconstruction constraint establishes a solid foundation for the success of ICA when the data is whitened.

## VIII. Conclusion

This paper presents OICNet, an online neural network designed for real-time EEG source separation using ICA. Our experimental results demonstrate that OICNet outperforms other methods in terms of computational complexity and convergence speed, highlighting its effectiveness and practicality for online source separation in diverse EEG applications. The superior performance of OICNet holds great promise for enhancing the accuracy and reliability of various realtime brain-computer interface (BCI) applications. Its online nature and computational efficiency make it particularly well-suited for real-time EEG-based BCI systems, where rapid and dependable performance is of utmost importance. Moreover, the identifiability of the extracted EEG features by OICNet is comparable to that of the offline standard Infomax ICA. By leveraging the strengths of OICNet, researchers and practitioners can benefit from its robust capabilities in real-time EEG source separation, leading to improved signal quality, enhanced feature extraction, and more accurate interpretation of brain activity. This, in turn, can facilitate the development of advanced BCI systems for a wide range of applications.

## REFERENCES

[1] Anthony J Bell and Terrence J Sejnowski. The “independent components” of natural scenes are edge filters. Vision research, 37(23):3327–3338, 1997.

[2] Pierre Comon. Independent component analysis, a new concept? Signal processing, 36(3):287–314, 1994.

[3] Scott Makeig, Anthony Bell, Tzyy-Ping Jung, and Terrence J Sejnowski. Independent component analysis of electroencephalographic data. Advances in neural information processing systems, 8, 1995.

[4] Scott Makeig, Tzyy-Ping Jung, Anthony J Bell, Dara Ghahremani, and Terrence J Sejnowski. Blind separation of auditory event-related brain responses into independent components. Proceedings of the National Academy of Sciences, 94(20):10979–10984, 1997.

[5] Christopher J James and Christian W Hesse. Independent component analysis for biomedical signals. Physiological measurement, 26(1):R15, 2004.

[6] Amar Kachenoura, Laurent Albera, Lotfi Senhadji, and Pierre Comon. Ica: a potential tool for bci systems. IEEE Signal Processing Magazine, 25(1):57–68, 2007.

[7] Ralph G Andrzejak, Klaus Lehnertz, Florian Mormann, Christoph Rieke, Peter David, and Christian E Elger. Indications of nonlinear deterministic and finite-dimensional structures in time series of brain electrical activity: Dependence on recording region and brain state. Physical Review E, 64(6):061907, 2001.

[8] Michal Teplan et al. Fundamentals of eeg measurement. Measurement science review, 2(2):1–11, 2002.

[9] Xiao Jiang, Gui-Bin Bian, and Zean Tian. Removal of artifacts from eeg signals: a review. Sensors, 19(5):987, 2019.

[10] Tzyy-Ping Jung, Scott Makeig, Marissa Westerfield, Jeanne Townsend, Eric Courchesne, and Terrence J Sejnowski. Analyzing and visualizing single-trial event-related potentials. Advances in neural information processing systems, 11, 1998.

[11] Tzyy-Ping Jung, Scott Makeig, Colin Humphries, Te-Won Lee, Martin J Mckeown, Vicente Iragui, and Terrence J Sejnowski. Removing electroencephalographic artifacts by blind source separation. Psychophysiology, 37(2):163–178, 2000.

[12] J-F Cardoso and Beate H Laheld. Equivariant adaptive source separation. IEEE Transactions on signal processing, 44(12):3017–3030, 1996.

[13] Shun-Ichi Amari. Natural gradient works efficiently in learning. Neural computation, 10(2):251–276, 1998.

[14] Sergio Cruces-Alvarez, Andrzej Cichocki, and Luis Castedo-Ribas. An iterative inversion approach to blind source separation. IEEE Transactions on Neural Networks, 11(6):1423–1437, 2000.

[15] Muhammad Tahir Akhtar, Tzyy-Ping Jung, Scott Makeig, and Gert Cauwenberghs. Recursive independent component analysis for online blind source separation. In 2012 IEEE International Symposium on Circuits and Systems (ISCAS), pages 2813–2816. IEEE, 2012.

[16] Jiantao Lu, Wei Cheng, and Yanyang Zi. Online blind source separation method with adaptive step size in both time-invariant and time-varying cases. Measurement Science and Technology, 31(4):045102, 2020.

[17] Sheng-Hsiou Hsu, Luca Pion-Tonachini, Tzyy-Ping Jung, and Gert Cauwenberghs. Tracking non-stationary eeg sources using adaptive online recursive independent component analysis. In 2015 37th Annual International Conference of the IEEE Engineering in Medicine and Biology Society (EMBC), pages 4106–4109. IEEE, 2015.

[18] Sheng-Hsiou Hsu, Tim R Mullen, Tzyy-Ping Jung, and Gert Cauwenberghs. Real-time adaptive eeg source separation using online recursive independent component analysis. IEEE transactions on neural systems and rehabilitation engineering, 24(3):309–319, 2015.

[19] Aapo Hyvarinen. Fast and robust fixed-point algorithms for independent component analysis. IEEE transactions on Neural Networks, 10(3):626–634, 1999.

[20] Te-Won Lee, Mark Girolami, and Terrence J Sejnowski. Independent component analysis using an extended infomax algorithm for mixed subgaussian and supergaussian sources. Neural computation, 11(2):417–441, 1999.

[21] Aapo Hyvärinen and Petteri Pajunen. Nonlinear independent component analysis: Existence and uniqueness results. Neural networks, 12(3):429–439, 1999.

[22] Philemon Brakel and Yoshua Bengio. Learning independent features with adversarial nets for non-linear ica. arXiv preprint arXiv:1710.05050, 2017.

[23] Hlynur DavíHlynsson and Laurenz Wiskott. Learning gradient-based ica by neurally estimating mutual information. In Joint German/Austrian Conference on Artificial Intelligence (Künstliche Intelligenz), pages 182–187. Springer, 2019.

[24] Hongming Li, Shujian Yu, and José C Príncipe. Deep deterministic independent component analysis for hyperspectral unmixing. In ICASSP 2022-2022 IEEE International Conference on Acoustics, Speech and Signal Processing (ICASSP), pages 3878–3882. IEEE, 2022.

[25] Anthony J Bell and Terrence J Sejnowski. An information-maximization approach to blind separation and blind deconvolution. Neural computation, 7(6):1129–1159, 1995.

[26] Luca Pion-Tonachini, Sheng-Hsiou Hsu, Scott Makeig, Tzyy-Ping Jung, and Gert Cauwenberghs. Real-time EEG source-mapping toolbox (REST): Online ICA and source localization. In 2015 37th Annual International Conference of the IEEE Engineering in Medicine and Biology Society (EMBC), pages 4114–4117. IEEE, 2015.

[27] Arnaud Delorme and Scott Makeig. Eeglab: an open source toolbox for analysis of single-trial eeg dynamics including independent component analysis. Journal of neuroscience methods, 134(1):9–21, 2004.

[28] Mohamed Ishmael Belghazi, Aristide Baratin, Sai Rajeswar, Sherjil Ozair, Yoshua Bengio, Aaron Courville, and R Devon Hjelm. Mine: mutual information neural estimation. arXiv preprint arXiv:1801.04062, 2018.

[29] Aapo Hyvärinen. New approximations of differential entropy for independent component analysis and projection pursuit. Advances in neural information processing systems, 10, 1997.

[30] Aapo Hyvärinen. Independent component analysis by minimization of mutual information. 1997.

[31] Aapo Hyvärinen and Erkki Oja. Independent component analysis: algorithms and applications. Neural networks, 13(4-5):411–430, 2000.

[32] Quoc Le, Alexandre Karpenko, Jiquan Ngiam, and Andrew Ng. ICA with reconstruction cost for efficient overcomplete feature learning. Advances in neural information processing systems, 24, 2011.

[33] J. Karhunen A. Hyvarinen and E. Oja. Independent Component Analysis. John Wiley Sons, Ltd.

[34] Xiaolong Zhu, Xianda Zhang, and Jimin Ye. Natural gradient-based recursive least-squares algorithm for adaptive blind source separation. Science in China Series F: Information Sciences, 47(1):55–65, 2004.

[35] Diederik P Kingma and Jimmy Ba. Adam: A method for stochastic optimization. arXiv preprint arXiv:1412.6980, 2014.

[36] Perrin Margaux, Maby Emmanuel, Daligault Sébastien, Bertrand Olivier, and Mattout Jérémie. Objective and subjective evaluation of online error correction during p300-based spelling. Advances in Human-Computer Interaction, 2012, 2012.

[37] Michael Falkenstein, Jörg Hoormann, Stefan Christ, and Joachim Hohns-bein. Erp components on reaction errors and their functional significance: a tutorial. Biological psychology, 51(2-3):87–107, 2000.

[38] maucle Wendy Kan Jérémie Mattout, Manu. Bci challenge @ ner 2015, 2014.

[39] Arnaud Delorme, Jason Palmer, Julie Onton, Robert Oostenveld, and Scott Makeig. Independent eeg sources are dipolar. PloS one, 7(2):e30135, 2012.

[40] Alexandre Gramfort, Martin Luessi, Eric Larson, Denis A Engemann, Daniel Strohmeier, Christian Brodbeck, Roman Goj, Mainak Jas, Teon Brooks, Lauri Parkkonen, et al. Meg and eeg data analysis with mne-python. Frontiers in neuroscience, page 267, 2013.

[41] Aapo Hyvärinen. Testing the ica mixing matrix based on inter-subject or inter-session consistency. NeuroImage, 58(1):122–136, 2011.

[42] Francesco Di Russo, Antíigona Martíinez, Martin I Sereno, Sabrina Pitzalis, and Steven A Hillyard. Cortical sources of the early components of the visual evoked potential. Human brain mapping, 15(2):95–111, 2002.

[43] Nitin Bansal, Xiaohan Chen, and Zhangyang Wang. Can we gain more from orthogonality regularizations in training deep networks? Advances in Neural Information Processing Systems, 31, 2018.

